# A gene-based capture assay for surveying patterns of genetic diversity and insecticide resistance in a worldwide group of invasive mosquitoes

**DOI:** 10.1101/2021.08.24.457535

**Authors:** Matthew L. Aardema, Michael G. Campana, Nicole E. Wagner, Francisco C. Ferreira, Dina M. Fonseca

## Abstract

Understanding patterns of diversification, genetic exchange, and pesticide resistance in insect species of human health concern is necessary for effective population reduction and management. With the broad availability of next-generation sequencing technologies, one of the best approaches for surveying such patterns involves the simultaneous genotyping of many samples for large numbers of genetic markers from across the known genome. To this end, the targeting of gene sequences of known function or inheritance can be a cost-effective strategy. One insect group of substantial health concern are the mosquito taxa that make up the *Culex pipiens* complex. Members of this complex transmit damaging arboviruses and filariae worms to humans, as well as other pathogens that are detrimental to endangered vertebrate species such as bird malaria. Here we describe our development of a targeted gene-based assay for surveying genetic diversity and population structure in this mosquito complex. To test the utility of this assay, we examined taxonomic divergence among samples from several members of the complex, as well as distinct populations of the relatively under-studied *Culex quinquefasciatus*, an urban pantropical species. We also examined the presence of known insecticide-resistance conferring alleles. Broadly, our developed gene-based assay proved effective for examining patterns of taxonomic and geographic clustering within the species complex, as well as for surveying genetic variants that have been associated with insecticide resistance. This assay will be useful for future studies that aim to understand the genetic mechanisms underlying the evolution of ubiquitous and increasingly damaging disease vectors.

## 1 INTRODUCTION

The brown dusk-biting mosquitoes collectively classified within the *Culex pipiens* complex (Diptera: Culicidae), include two globally distributed invasive species, the temperate *Culex pipiens*, and the tropical *Cx. quinquefasciatus*, along with several additional taxa with more restricted distributions (reviewed in Aardema et al. 2021). Specific populations of these two species are critical urban vectors of human periodic filariasis (*Wuchereria bancrofti*) and several epidemic encephalitides such as West Nile virus (Kramer et al. 2008) and Usutu virus (Eiden et al. 2018). They also vector avian malaria, a group of parasites that have become of critical concern to island bird communities in Hawaii, the Galapagos, and elsewhere (Bataille et al. 2009, McClure et al. 2020, Paxton et al. 2016).

The current distribution of many cosmopolitan mosquito species was likely driven by rapid human movement, and accordingly these distributions are a relatively recent phenomenon (Gippet et al. 2019). One of the best-known invasive species is the yellow fever mosquito, *Aedes aegypti*. Outside its source location in Africa, populations of *Ae. aegypti* all share the same basic genotype, which is evidence of its rapid expansion associated with human movement (Gloria-Soria et al. 2016). Interestingly, in contrast to this pattern, microsatellite analyses of populations of *Cx. pipiens* and *Cx. quinquefasciatus* from across the world uncovered unexpectedly high levels of genetic diversity. For example, continental populations of *Cx. quinquefasciatus* flanking the Pacific Ocean are highly differentiated (Fonseca et al. 2006). Furthermore, although historical records pinpoint an original introduction of *Cx. quinquefasciatus* into the Hawaiian Islands from the Americas (Dine 1904), current Hawaiian *Cx. quinquefasciatus* have a distinct Australasian signature. The mechanisms underlying the replacement of the first strain by the second are unknown and understanding them will likely require a better understanding of the specific genetic makeup (i.e. which genes and their capabilities) of the population(s) involved.

Another interesting phylogenetic phenomenon observed within the *Cx. pipiens* complex is that patterns of genetic divergence are not always associated with obvious phenotypic differences. For example, while it was found that the two described ecological forms of *Cx. pipiens* (which differ in behavior and physiology) have diagnostic multilocus genotypes that distinguish them (Fonseca et al. 2004), this analysis uncovered the existence of genetic exchange between the two forms. Such inter-taxonomic hybridization may have potentially significant negative consequences for arboviral transmission to humans. Several studies have also found evidence of extensive hybrid zones between temperate *Cx. pipiens* or *Cx. pipiens pallens* (a subspecies limited to northeastern Asia) and tropical *Cx. quinquefasciatus* (Fonseca et al. 2009, Kothera et al. 2009). Analysis of genetic variation at the acetylcholinesterase locus 2 (ACE2) across members of the complex indicated that the hybridization event that resulted in the temperate *Cx. pipiens pallens* was unidirectional which is surprising since current patterns of hybridization with *Cx. quinquefasciatus* are by-directional (Fonseca et al. 2009).

Next-generation sequencing (NGS) has enabled vast amounts of data to be collected at relatively low cost (e.g. Goodwin et al. 2016, Kulkarni and Frommolt 2017). However, mosquito genomes are often riddled with repetitive DNA, making whole genome data collection and analysis expensive and wasteful since only a small proportion of the genetic variation observed can be confidently compared across all specimens. Challenges for data collection and analysis are also created by the presence of diverse microbial symbionts such as *Wolbachia* and endogenous viral elements (e.g. Whitfield et al. 2017). Capitalizing on recent technological advancements, a capture approach where DNA or RNA probes designed to match known genes are hybridized to DNA libraries of individual specimens and sequenced has been gaining traction (e.g. Campana et al. 2016, Cassin-Sackett et al. 2019, Quek et al. 2020). Because it bypasses large amounts of DNA of unknown function and heritability, a gene targeted approach allows users to pool tens or even hundreds of indexed specimens, and cost-effectively uncover thousands of homologous loci simultaneously. However, such methodologies have so far had minimal use for examining population genetics or evolutionary patterns in mosquitos (but see Itokawa et al. 2019).

Here we describe the design and use of a genetic baits assay targeting 512 genes annotated in the *Cx. quinquefasciatus* genome. Among these genes are a large number of known regions that have been shown to harbor genetic variation that correlates with insecticide resistance. We examined the utility of these baits for taxonomic differentiation and patterns of admixture by sequencing samples from four taxa of the *Cx. pipiens* species complex, samples of known hybrid origin, and one sample of a closely related, outgroup taxon, *Culex torrentium*. To further examine the potential of these baits for exploring finer scale, intra-taxonomic population structure and differentiation, we included samples of *Cx. quinquefasciatus* from multiple geographic sources. Finally, we investigated the presence and frequency of known insecticide resistance alleles among the samples examined.

## 2 MATERIALS AND METHODS

### 2.1 Bait design and screening

We designed an in-solution capture assay targeting 131 rapidly evolving *Culex* genes obtained from a comparison of *de novo* whole transcriptomes from *Cx. pipiens* f. pipiens and f. molestus (Price & Fonseca, 2015) using the *Cx. quinquefasciatus* genome (CpipJ1.3 Johannesburg, South Africa (Arensburger et al. 2010) as a reference (Table S1). Of the seven enriched GO terms identified, five terms (chitin metabolic process, chitin binding, serine-type endopeptidase activity, proteolysis and odorant binding) were also enriched along the ‘fly’ branch (Adams et al., 2000) indicating they may represent a genetic ‘core’ for adaptive evolution within the Diptera. To estimate genotyping error rates, we included 28 slow-evolving genes (Price & Fonseca, 2015). We designed baits to cover the complete exonic and intronic regions for simultaneous investigation of both adaptive and neutral evolution. We also added capture probes for 353 genes involved in insecticide resistance: P450s, alpha and beta esterases, sodium channel, and acetylcholinesterase (Asgharian et al., 2015). To ensure optimal capture, Daicel Arbor Biosciences designed 39,953 120 bp baits with ∼1.5x flexible tiling density (∼80bp probe spacing) to cover these targets. Bait candidates were accepted when they satisfied one of the following conditions: a) no blast hit with a T_m_ above 60°C, b) no more than two hits at 62.5 - 65°C, or 10 hits in the same interval and at least one neighbor candidate being rejected. c) no more than 2 hits at 65 - 67.5°C and 10 hits at 62.5 - 65°C and two neighbor candidates on at least one side being rejected. d) no more than a single hit at or above 70C or e) no more than one hit at 65 - 67.5°C and 2 hits at 62.5 - 65°C and two neighbor candidates on at least one side being rejected. The baits were synthesized as a myBaits version 3 kit. After this “stringent” filtration, 29,992 baits were retained, covering all 512 target genes with at least one bait and 2,524,269 bp of the targeted sequence.

### 2.2 Target enrichment and sample sequencing

We chose specimens representative of the genetic diversity observed across the complex (Table 2). Specifically, we analyzed specimens of the two *Culex pipiens* forms from Europe and North America, specimens of the subspecies *Cx. pipiens pallens* from the Republic of Korea and, to assess the power of the assay to discern intra-specific patterns, also specimens of *Cx. quinquefasciatus* from six distinct geographic regions: east-southeast Asia, Samoa, Hawaii, North America (including the Caribbean), Brazil and Nigeria. We also included known hybrids of *Cx. pipiens* and *Cx. quinquefasciatus* from California and North Carolina. Most specimens had already been examined using a panel of microsatellite loci (Fonseca et al. 2004, 2006, 2009, Strickman & Fonseca 2012).

We extracted DNA from individual mosquitoes using a phenol-chloroform method detailed in Fonseca et al (2000). We then performed an initial step to clean and concentrate DNA by using Omega Mag-Bind^®^ TotalPure NGS beads at 0.9 ratio following manufacturer’s protocol. For library preparation, we used Illumina^®^ DNA library prep (formerly Nextera DNA Flex) following the manufacturer’s protocol. Concentration and quality of the libraries were measured using Qubit^®^ 2.0 Fluorometer and Bioanalyzer High Sensitivity DNA Analysis kit (Agilent), respectively. To create amplicons that do not have affinity to streptavidin, we performed four amplification cycles following instructions in Appendix A2 of myBaits Hybridization Capture for NGS protocol (v. 4.01). For that, we used universal P5 and P7 primers. These products were cleaned using Omega Mag-Bind^®^ beads and hybridized with our capture biotinylated baits for target enrichment following myBaits protocol (v. 4.01). We used diluted baits to a ratio of 1:6. These libraries were amplified following 12 cycles using KAPA^®^ HiFi Hotstart ready mix, and resulting products were cleaned with AMPure XP beads (Beckman Coulter). Concentration and quality of final libraries were checked using Qubit^®^ and Bioanalyzer, and each sample was adjusted to a final concentration of 4 nM (1.33 ng/µl). We obtained libraries with fragment sizes of 600 bp in average. These were 2 × 300 bp paired-end sequenced in groups of 6 or 7 on an Illumina MiSeq using 600-cycle Miseq version 3 kits.

### 2.3 Data mapping and variant calling

We first used the program Trim Galore (https://github.com/FelixKrueger/TrimGalore) to trim Illumina sequencing adapters and bases from read ends with a quality score less than 20. We then removed both reads of a pair if either read was less than 30 bases long. Next, we mapped the remaining trimmed reads to the *Cx. quinquefasciatus* reference genome (v. CpipJ2.5; Arensburger et al. 2010) which we obtained from VectorBase (https://vectorbase.org/vectorbase/app). Our mapping was done using the program BWA-MEM v. 0.7.12 with default settings (Li 2013). Next, we added read groups and sorted the mapped reads using the AddOrReplaceReadGroups function in Picard v. 1.119 (http://broadinstitute.github.io/picard/). We then marked read duplicates using the tool MarkDuplicates, also with Picard v. 1.119, followed by indel realignment using IndelRealigner in the Genome Analysis Toolkit (‘GATK’) v. 3.6 (McKenna et al. 2010). Finally, for each sample, we identified genetic variants using GATK’s HaplotypeCaller (specific flags: -- emitRefConfidence GVCF, --variant_index_type LINEAR, --variant_index_parameter 128000 - rf BadCigar).

With the resulting raw VCF files (one per sample), we used GATK’s GenotypeGVCFs function to produce a single, multi-sample VCF containing all identified variants observed across all samples. This file was then filtered to retain only single nucleotide polymorphisms (SNPs), using the SelectVariants tool in GATK v. 4.0.8.1. This tool was also used to remove any variants that fell outside our designated baits coordinates (Table S1). Next, we applied a series of hard quality filters, removing all SNPs with any of the following parameters: QD < 11.0, FS > 40.0, MQ < 56.0, MQRankSum < -0.2, ReadPosRankSum < -3.0, and/or SOR > 2.0. These thresholds were based on the observed distribution of variants (Figure S1), and were equal to, or stricter than, the recommended values given in GATK’s best practices. Finally, we used the program SnpEff v. 4.3 (Cingolani et al. 2012), with a custom database to annotate the remaining SNPs for downstream sorting by variant type.

### 2.4 Genetic clustering and admixture

We first wanted to examine genetic relationships and potential gene flow (admixture) across all the samples derived from the *Cx. pipiens* species complex. We also wanted to examine just the *Cx. quinquefasciatus* samples, to assess the utility of these baits for surveying intraspecific population relationships. To carry out these analyses, we wanted to maximize the number of selectively neutral markers available. Prior work showed the importance of utilizing a large number of segregating markers to detect clustering in genetic data when divergence between distinct populations is likely to be low (Patterson et al. 2006). For likely ‘neutral’ genetic markers, we used all variants that were annotated as either synonymous or intronic. Although work in *Drosophila* suggests that mutations in both of these site categories can experience selection (e.g. Shields et al. 1988, Halligan et al. 2004, Andolfatto 2005), the strength of this selection is likely far less than that acting on non-synonymous variation.

From our VCF database of high quality synonymous and intronic SNPS, we used GATK’s ‘SelectVariants’ tool to generate two new VCFs, one with all samples except the outgroup *Cx. torrentium* (henceforth ‘*Cx. pipiens* complex’ dataset), and a second with only the *Cx. quinquefasciatus* samples (henceforth ‘*Cx. quinquefasciatus*’ dataset). We then removed any variant from both datasets that was not in Hardy-Weinberg equilibrium (p < 0.0001), and any variant in which the minor allele was represented at less than 5% frequency. Both filtering steps were carried out using VCFtools v. 0.1.17 (Danecek et al. 2011). Finally, for both datasets, we used PLINK v.1.90b6.6 (Purcell et al. 2007) to remove SNPs with a pairwise squared correlation (r^2^) greater than 50% within sliding windows of 50 SNPs at 10 SNP increments between windows (Novembre et al. 2008). This was done to reduce the impact of linkage between SNPs on our examinations of population clustering and admixture (Pritchard et al. 2000).

We first used principal component analyses (PCAs) to investigate clustering among the samples in both datasets. These PCAs were conducted with the program PLINK v. 1.90b6.6 (Purcell et al. 2007), and the results visualized using R v. 4.0.2 (R Core Team 2020), focusing on the first two principal components (PC1 & PC2). We also examined patterns of genetic structure within our data using a maximum likelihood approach with the program ADMIXTURE v. 1.3.0 (Alexander et al. 2009). With ADMIXTURE we examined potential clusters (K) from one to seven in both datasets. Each K value was run 20 independent times with a different seed value for each run. Across K values, we compared the means observed for the standard error of the 10-fold cross-validation (CV) error estimate to determine the number of clusters best supported by the data (Alexander et al. 2015). We determined the average q-matrix cluster assignment for each sample for each K value using the online version of CLUMPAK (Kopelman et al. 2015), with default settings.

### 2.5 Genetic diversity and taxonomic divergence

To examine the amount of genetic diversity harbored within individual samples, populations and taxa, we used GATK v. 4.0.8.1 to designate all sample-variant combinations with a depth of coverage less than 15X to ‘no call’ status. A minimum read depth of 15X or greater was shown to be adequate for assessing the diploid state of an allele (homozygous vs. heterozygous) within samples (Song et al. 2016). Next, we used GATK to retain only biallelic SNPs that were annotated as either ‘synonymous’ or ‘intronic’ and called in all samples. This variant filtering was done to improve the equivalency of relative diversity estimates across all the samples. Finally, we used VCFtools v. 0.1.17 to count the number of observed homozygous variants. This was then used to calculate the average heterozygosity within a sample across assessed sites (Nei and Li 1979, Nei 1987). Taxon and population (*Cx. quinquefasciatus* only) means and standard errors of these means were then calculated. Although these estimates were not able to give us absolute estimates of genetic diversity (because they only included known segregating sites), they did allow us to make relative comparisons between groups of samples (e.g. taxa or populations).

To examine relative divergence between sample clusters (e.g. taxa or populations), we used VCFtools v. 0.1.17 and our larger clustering dataset to calculate the pairwise fixation index (F_st_, Weir and Cockerham 1984). Comparisons were done between the four complex taxa excluding the known hybrids, and between these and the outgroup *Cx. torrentium*. We also compared the *Cx. quinquefasciatus* population from the six designated geographic regions. All sample taxonomic and population designations were based on their prior assignments (Table S2). We report both the weighted and unweighted estimates. Weighted estimates may be more strongly impacted by unequal samples sizes, whereas unweighted estimates may be more affected by variants segregating at low frequencies (Weir 2002).

### 2.6 Phylogenetic analysis

To further examine sample clustering as well as taxonomic relationships amongst all samples, including the outgroup *Cx. torrentium*, we performed a maximum likelihood phylogenetic analysis. To do this, we wanted to utilize neutral variants that likely have similar mutation probabilities. Therefore, from our annotated variants dataset, we used BCFtools v. 1.9 (Li 2011) to select only 4-fold (‘silent’) segregating sites. Next, we removed variants that were not in Hardy-Weinberg equilibrium using VCFtools v. 0.1.17. We also thinned highly correlated SNPs as described above. The resulting VCF file was converted to PHYLIP format using the vcf2phylip.py v. 1.5 python script (https://zenodo.org/record/1257058#.YJL3ymZKi6t). We then used jModelTest 2.1.10 with default settings to select the best-fit model of nucleotide substitution for our datasets based on the BIC scores (Guindon and Gascuel 2003, Darriba et al. 2012). With the best fit model, we used PhyML v. 3.1 (Guindon et al. 2010) to carry out a maximum-likelihood phylogenetic analysis, with 100 non-parametric bootstrap replicates to determine confidence values. The resulting phylogenetic tree was visualized using the program FigTree v. 1.4.4 (Rambaut 2018).

### 2.7 Presence of variants potentially conferring insecticide resistance

To assess the utility of our capture assay for surveying known genetic variation that may contribute to insecticide resistance, we first conducted a literature survey to identify known single nucleotide variation that has been implicated in insecticide resistance within the mosquitoes of the *Cx. pipiens* complex. In particular, we examined published data which indicated the gene and position of a segregating variant. These were exclusively missense mutations that changed the amino acid sequence and likely protein interactions with the insecticide. With these genome coordinates (chromosome and base position), we used VCFtools v. 0.1.17 to calculate the frequencies of the susceptible and resistant alleles across the samples. We also used VCFtools to examine the sample-specific presence of these variants to compare taxa and populations.

## 3 RESULTS

### 3.1 Data mapping and variant calling

The number of raw read pairs, read pairs after filtering, and the number of properly mapped reads for each sample are given in Table S3. Also given in this table are the percentage of reads mapped and the average variant depth for all samples after filtering.

We initially called 12,301,010 variants across all samples, including both single nucleotide polymorphisms (SNPs) and insertions/deletions (INDELs). After removing all INDELs and any additional variants not located in our designated baits, we were left with 315,512 SNPs. Quality filtering further reduced this to 132,185 SNPs.

### 3.2 Genetic clustering and admixture

For examining genetic relationships for all the samples within the *Culex pipiens* complex, we generated a dataset consisting of SNPs annotated as either ‘synonymous’ or ‘intronic’ and that were in Hardy-Weinberg equilibrium. This dataset contained 14,303 unlinked variants.

Our principal component analysis with this dataset revealed the greatest genetic divergence (PC1) occurs between the samples designated as *Cx. quinquefasciatus* and all the other samples (Figure 1). PC2 distinguished the *Cx. pipiens pallens* samples from the other samples. As predicted, the two samples known to be admixed between *Cx. quinquefasciatus* and *Cx. pipiens* fell intermediate between these taxa along PC1. Additionally, along PC2, there appears a small distinction between the two forms of *Cx. pipiens* (f. *pipiens* and f. *molestus*), suggesting possible genetic divergence.

**Figure 1.**
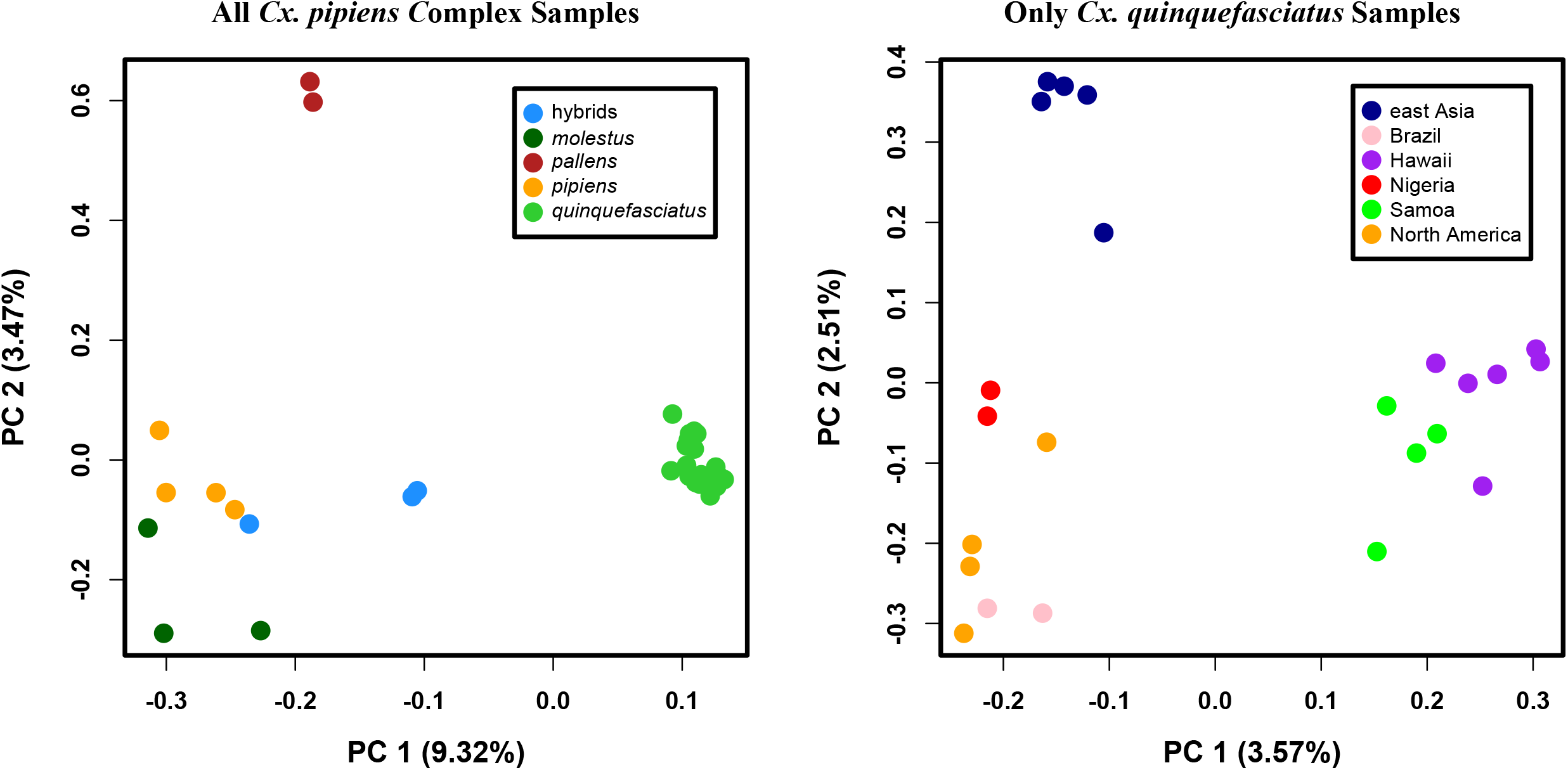
Results of Principal Component Analyses. Shown are the first and second principal components (PC1 & PC2) for all the *Cx. pipiens* complex samples (left panel) b) just the *Cx. quinquefasciatus* samples (right panel). These analyses were performed with neutral, segregating variants. Taxonomic and population memberships were based on prior designations.

The analysis of ADMIXTURE results with this dataset indicated that a K value of 2 was best supported (Figure S2). Population clustering at this K value again indicates the genetic distinction between the *Cx. quinquefasciatus* samples and the other complex samples (Figure 2). The two samples known to be hybrids between *Cx. pipiens* and *Cx. quinquefasciatus* clearly show this unique ancestry. At K=3 we saw a division between *Cx. quinquefasciatus* samples from Hawaii and Samoa and all other *Cx. quinquefasciatus* samples. At K=4 the *Cx. quinquefasciatus* samples were further subdivided. At K=5, the *Cx. pipiens pallens* samples were distinguished. Larger K values (6&7) further divide the *Cx. quinquefasciatus* samples and revealed samples with varying degrees of admixture.

**Figure 2.**
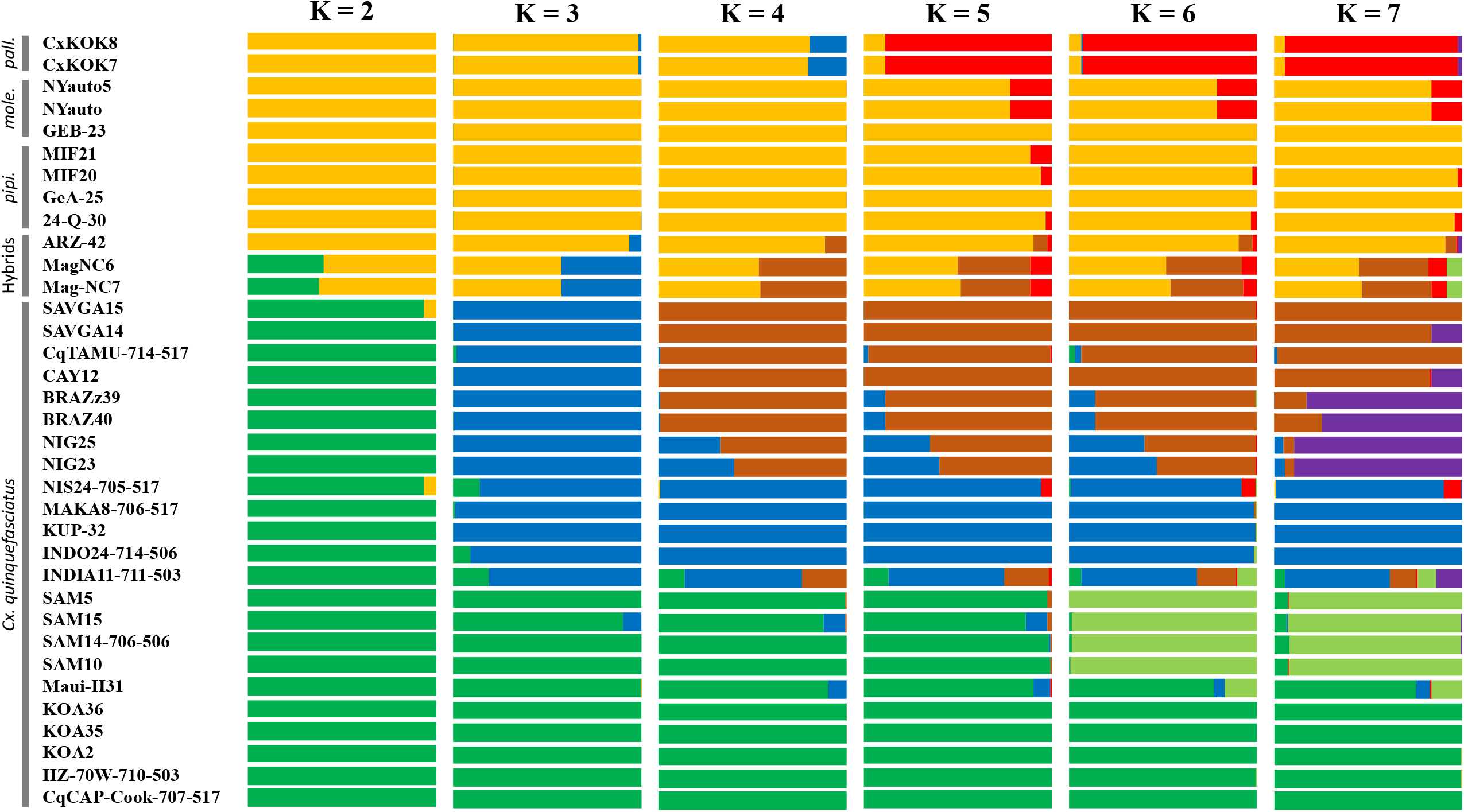
All complex ADMIXTURE results. Shown are the percent ancestry assignments (Q) for at K values 2 through 7 based on our analysis of admixture. Sample designations are given on the left along with taxonomic designations.

We also looked at sample clustering just in our known *Cx. quinquefasciatus* samples. This dataset consisted of 9,829 unlinked, segregating variants annotated as synonymous or intronic. All samples within the complex roughly clustered within their known geographic region (Figure 1), and more broadly there were three distinct groupings. These corresponded to a cluster of Hawaiian and Samoan samples that were distinct from all the other samples along PC1, and a cluster of east Asian samples that were distinct from the third cluster along PC2. This third cluster consisted of samples from North America and the Caribbean, Brazil and Nigeria.

Assessment of our admixture results for the *Cx. quinquefasciatus* specimens suggested a single taxonomic group (i.e. K=1; Figure S2). This is not surprising given the limited number of markers utilized and the low amounts of genetic divergence likely to be present between populations of this species (Patterson et al. 2006). The greatest distinction between populations within our *Cx. quinquefasciatus* samples was between those from Hawaii and Samoa, and the remaining samples (Figure 3). At K=3 we see the east Asian samples from a distinct cluster, recapitulating the results for our principal component analysis. One sample, from India (INDIA11-711-503) appears to be highly admixed with genetic representation from multiple populations across K values. At K=4, the Hawaiian and Samoan samples form distinct clusters. The Nigerian and Brazilian samples show their distinctiveness (and relation to one another) at K=5. However, this affiliation disappears at K=6. Such cluster shifting across K values highlights the overall degree of genetic similarity among these samples and potentially reveal the limitations of this dataset for examining fine-scale structuring between closely related populations.

**Figure 3.**
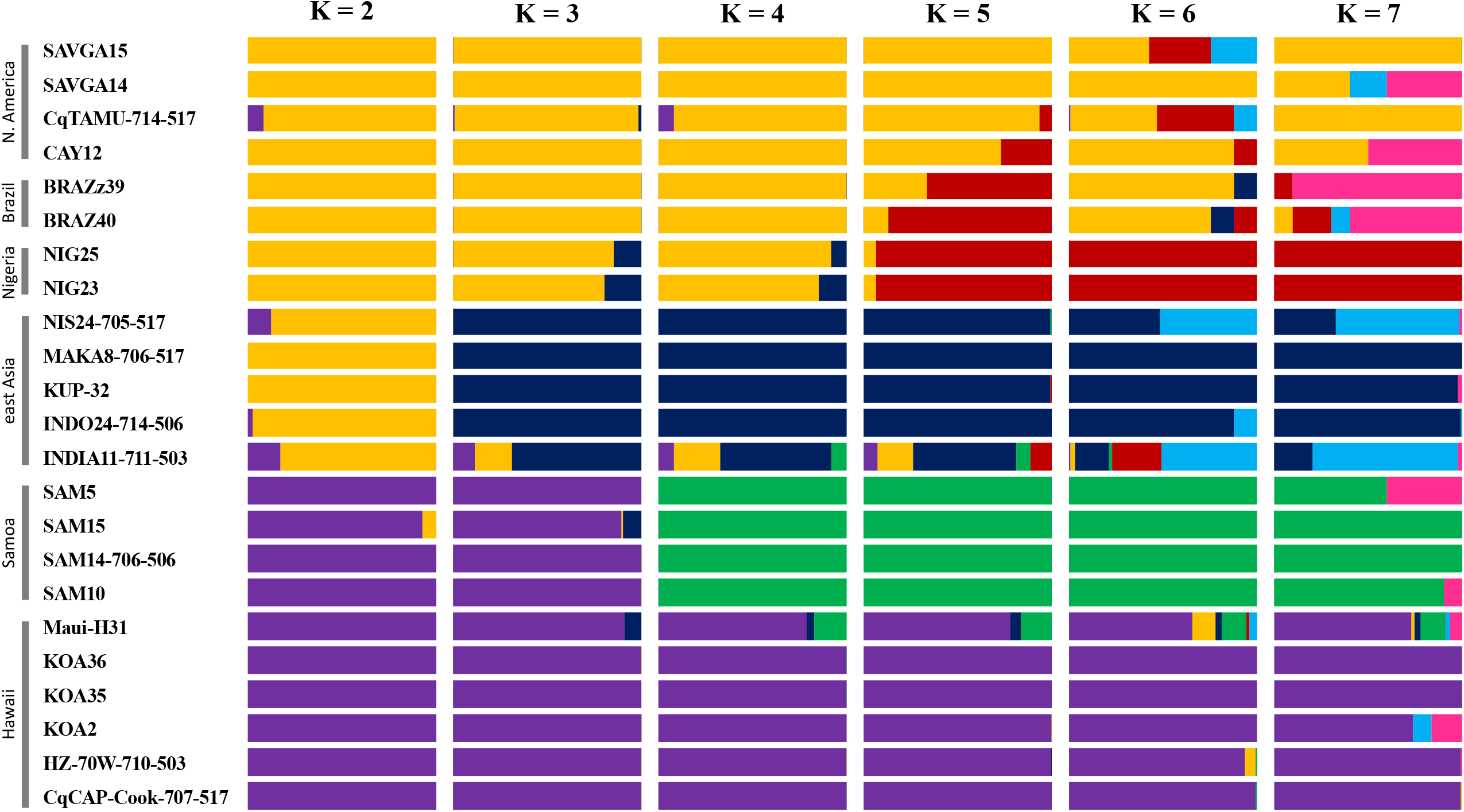
*Cx. quinquefasciatus* ADMIXTURE results. Shown are the percent ancestry assignments (Q) for at K values 2 through 7 based on our analysis of admixture. Sample designations are given on the left along with population designations.

### 3.3 Genetic Diversity and Taxonomic Divergence

To examine relative genetic diversity within these mosquitoes, we utilized 916 biallelic, neutral SNPs which each had a depth of at least 15 reads (15X) in all samples. The mean number of heterozygous sites and the mean sample pairwise heterozygosity (π) for all taxa are given in Table 1, and each sample’s individual diversity observations are given in Table S4. Surprisingly the taxon/group with the highest π values was *Cx. pipiens pallens* at 0.091 (SE=0.006). This value means that among the *Cx. pipiens pallens*, on average 9.1% of the 916 SNPs were found in a heterozygous state. The next highest value of π was observed in the *Cx. torrentium* sample with 0.084. The known hybrids had an average π of 0.066 (SE=0.009). The lowest mean π value was observed in the *Cx. quinquefasciatus* samples (0.023, SE=0.002).

**Table 1.** Relative genetic diversity within taxa across 916 neutral, bi-allelic, segregating SNPs. Given are the taxonomic designations (including a category for known hybrid samples), sample size for each taxon, the mean number of heterozygous sites observed per sample with standard error, and the corresponding mean pairwise sample heterozygosity value with standard error.

| **Taxon** | **Sample Size (n)** | **Mean Number of Heterozygous Sites (SE)** | **Mean Sample Pairwise Heterozygosity (SE)** |
| --- | --- | --- | --- |
| Known Hybrids | 3 | 60.7 (7.9) | 0.066 (0.009) |
| *Cx. pipiens* f. *molestus* | 3 | 25.0 (11.0) | 0.027 (0.012) |
| *Cx. pipiens* f. *pipiens* | 4 | 62.3 (7.2) | 0.068 (0.008) |
| *Cx. pipiens pallens* | 2 | 83.5 (5.5) | 0.091 (0.006) |
| *Cx. quinquefasciatus* | 23 | 20.9 (2.1) | 0.023 (0.002) |
| *Cx. torrentium* | 1 | 77 | 0.084 |

To examine relative genetic diversity within just the *Cx. quinquefasciatus* samples, we utilized 540 SNPs that were determined to be biallelic and had a depth of at least 15 reads (15X) in the samples under consideration. Furthermore, these were deemed neutral by virtue of being annotated as ‘synonymous’ or ‘intronic’. The mean number of heterozygous sites and the mean sample pairwise heterozygosity (π) for the six geographic designations of *Cx. quinquefasciatus* are given in Table 2. The samples from east Asia has the highest mean observed π with a value of 0.150 (SE=0.015). Hawaiian samples also appeared to be relatively genetically diverse with a π value of 0.103 (SE=0.017). The lowest mean values of π were observed in the Samoan (0.070, SE=0.013) and Brazilian samples (0.019, SE=0.012).

**Table 2.** Relative genetic diversity within populations of *Cx. quinquefasciatus* across 540 segregating, neutral, bi-allelic SNPs. Given are the population designation, sample size for each population, the mean number of heterozygous sites observed per sample with standard error, and the corresponding mean pairwise sample heterozygosity value with standard error.

| **Population** | **Sample Size**<br>**(n)** | **Mean Number**<br>**of**<br>**Heterozygous**<br>**Sites** | **Mean Sample**<br>**Pairwise**<br>**Heterozygosity** |
| --- | --- | --- | --- |
| east Asia | 5 | 81.0 (7.9) | 0.150 (0.015) |
| Samoa | 4 | 38.0 (6.9) | 0.070 (0.013) |
| Hawaii | 6 | 55.5 (8.9) | 0.103 (0.017) |
| North America &<br>Caribbean | 4 | 46.5 (8.5) | 0.086 (0.016) |
| Brazil | 2 | 10.5 (6.5) | 0.019 (0.012) |
| Nigeria | 2 | 48.0 (11.0) | 0.089 (0.020) |

Table 3 give the pairwise unweighted and weighted estimates of the fixation index (F_st_, Weir and Cockerham 1984), between each of the four *Cx. pipiens* complex taxa examined here as well as the outgroup, *Cx. torrentium*. Weighted estimates were always larger than unweighted estimates. Not surprisingly, the highest values were observed between the complex taxa and the *Cx. torrentium* sample. Among the taxa within the *Cx. pipiens* complex, the highest unweighted F_st_ value was between *Cx. quinquefasciatus* and *Cx. pipiens* f. *pipiens* (0.2967). With weighted F_st_ values, the highest was between *Cx. quinquefasciatus* and *Cx. pipiens* f. *molestus* (0.6415). The lowest estimated values were between the two *Cx. pipiens* forms (unweighted= -0.1026, weighted= 0.0276).

**Table 3.**
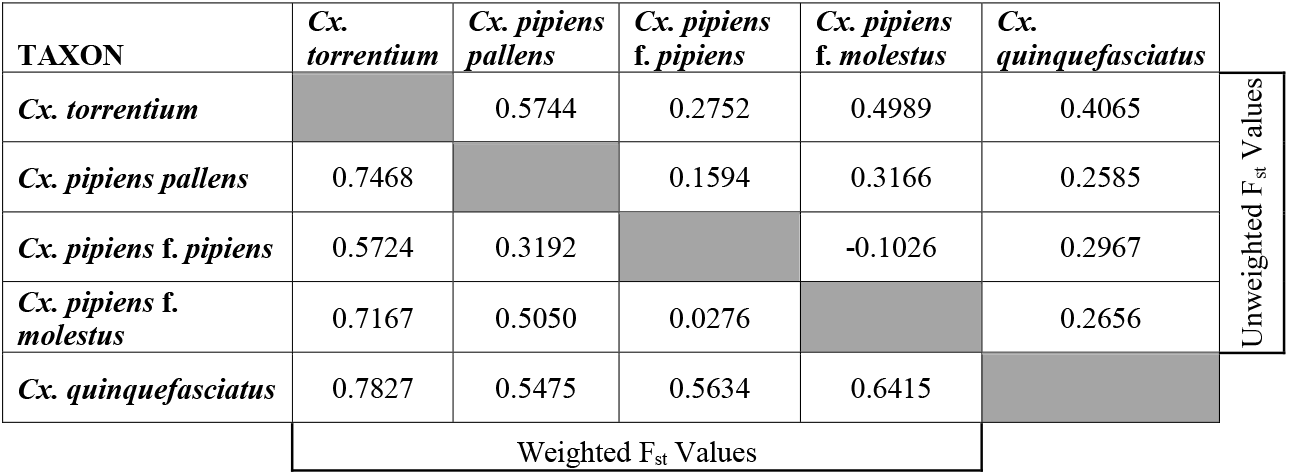
Pairwise F_st_ values between taxa. Given are both unweighted and weighted measures for each pair of taxa (excluding known hybrid samples). Taxonomic designations were designated prior to this study (see text for more details).

Pairwise unweighted and weighted estimates of F_st_ between the six designated geographic populations of *Cx. quinquefasciatus* are given in Table 4. Again, the weighted estimates were always larger than the unweighted estimates. For both estimate types, the highest values were observed between samples from Nigeria and Samoa (unweighted= 0.1233, weighted= 0.2387). For unweighted F_st_ values, the lowest estimate was between samples from Brazil and North America, including the Caribbean (−0.2329). The lowest estimated weighted F_st_ value was between Brazilian and east Asian samples (−0.0668).

**Table 4.** Pairwise F_st_ values between *Cx. quinquefasciatus* populations. Given are both unweighted and weighted measures for each pair of populations. Population designations were determined based on collection location (see text for more details).

| POPULATION | east Asia | Samoa | Hawaii | North America & Caribbean | Brazil | Nigeria | Unweighted $F_{st}$ Values |
| --- | --- | --- | --- | --- | --- | --- | --- |
| east Asia |  | 0.0514 | 0.0593 | 0.0051 | -0.2280 | 0.0161 |  |
| Samoa | 0.1611 |  | 0.0168 | 0.1016 | -0.0514 | 0.1233 |  |
| Hawaii | 0.1326 | 0.1026 |  | -0.0265 | -0.1741 | 0.0578 |  |
| North America & Caribbean | 0.1104 | 0.2277 | 0.0327 |  | -0.2329 | 0.0342 |  |
| Brazil | -0.0668 | 0.1622 | 0.0234 | -0.0638 |  | -0.1288 |  |
| Nigeria | 0.1100 | 0.2387 | 0.1626 | 0.1389 | 0.0497 |  |  |
| Weighted $F_{st}$ Values | | | | | | | |

### 3.4 Phylogenetic analysis

Our dataset for phylogenetic analysis consisted of 1,735 unlinked 4-fold synonymous SNPs, all of which were present in at least 75% of the samples. The evaluation of models of nucleotide sequence evolution indicated that a transversional model of mutation with a gamma distribution of rate heterogeneity best fit the data (TVM + Γ, Tavaré 1986). As expected, the outgroup species, *Cx. torrentium*, was clearly distinct from the samples of the *Cx. pipiens* complex (Figure 4). The *Cx. quinquefasciatus* samples also clustered with high confidence and overall the clustering recapitulated the results of the PCAs and Structure analyses.

**Figure 4.**
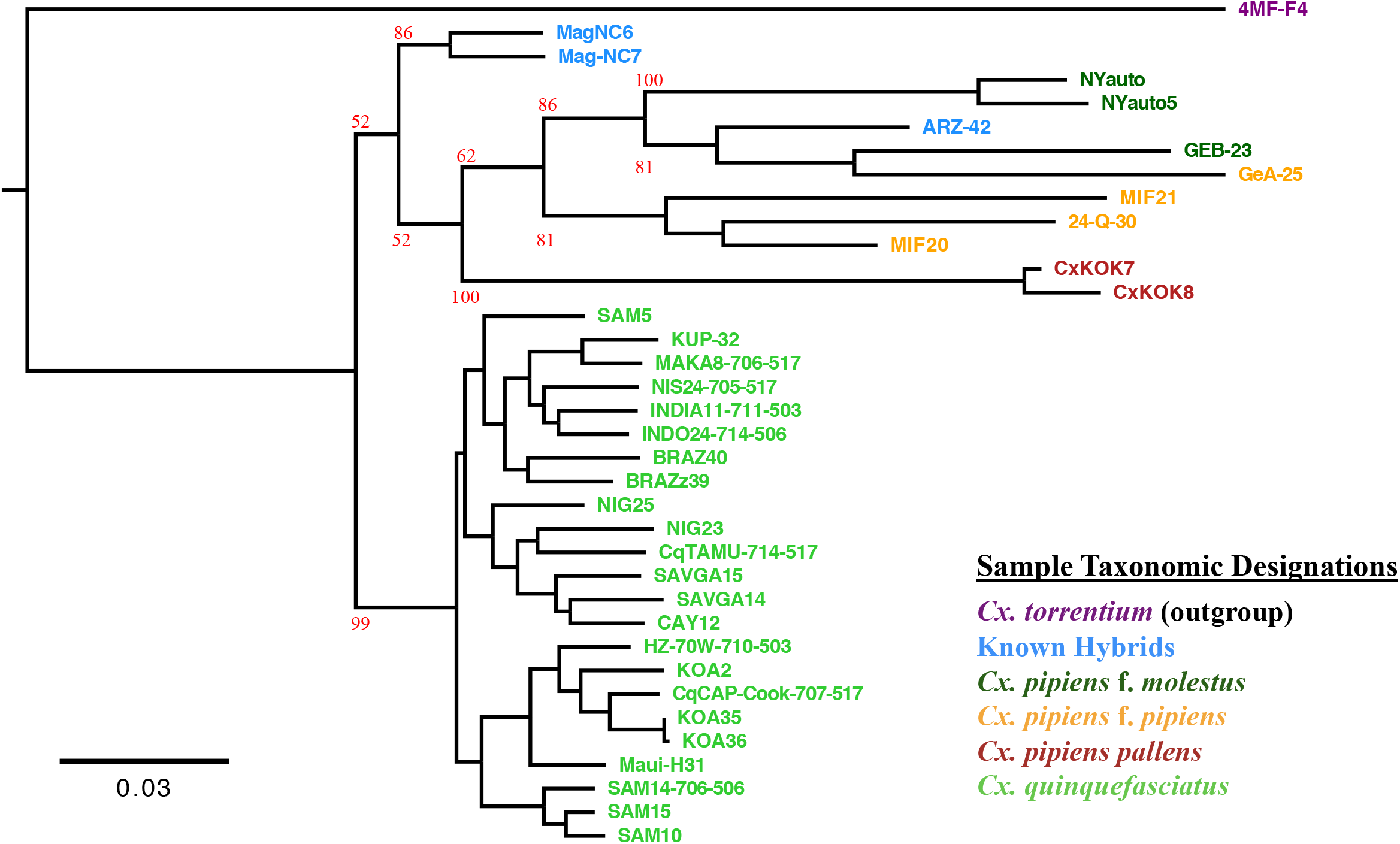
Maximum-likelihood phylogeny of samples. A maximum-likelihood analysis of all samples using a transversional model of mutation with a gamma distribution of rate heterogeneity. 100 bootstrap replicates of the analysis were performed and the bootstrap support for major nodes are shown in red. The colors correspond to the different taxonomic designations.

### 3.5 Presence of variants potentially conferring insecticide resistance

From our literature review of genetic variation found to confer insecticide resistance in the *Cx. pipiens* complex, we investigated the presence and frequency of seven single nucleotide polymorphisms among our samples (Table 5). All resistance-conferring alleles were present in the samples we examined here. For one of these sites, R213, found in gene CYP6BZ2 (cytochrome P450 6BZ2), there are two allelic changes that may confer resistance (R213L and R213Q). No sample had a copy of both resistance-conferring alleles. However, only four samples were homozygous for the susceptible nucleotides at this site. Of the eight possible resistance-conferring alleles at seven sites, only those in the cytochrome P450, 6BZ2 gene were observed at frequencies greater than 20% (T:41.7%, A:33.3%) across all surveyed mosquito samples. All other resistance alleles were found at lower frequencies than their alternative, susceptible allele.

**Table 5.** Summary of Insecticide Resistance Allele Frequencies. Given are the genomic position of examined resistance-conferring SNPS (chromosome and base position), gene ID (in the annotated *Cx. quinquefasciatus* genome), gene name, amino acid change, number of chromosomes examined (# of samples the variant is called at position * 2), and the frequencies of the susceptible and resistant alleles.

| Chromosome | Position | Gene ID | Gene Name | Amino Acid Change | Number of Chromosomes Examined (n=72) | Susceptible Allele Frequency | Resistance Allele Frequency | Reference |
| --- | --- | --- | --- | --- | --- | --- | --- | --- |
| supercont3.35 | 864866 | CPIJ002538 | CYP6AG12: cytochrome P450 6AG12 | H293L | 72 | A:0.986 | T:0.014 | Kothera et al. 2019 |
| supercont3.104 | 245887 | CPIJ005956 | CYP6BZ2: cytochrome P450 6BZ2 | R213L, R213Q | 72 | G:0.250 | T:0.417, A:0.333 | Kothera et al. 2019 |
| supercont3.106 | 33980 | CPIJ006034 | ACE-1: acetylcholinesterase | G247S | 56 | G:0.929 | A:0.071 | Zhao et al. 2014 |
| supercont3.106 | 46973 | CPIJ006034 | ACE-1: acetylcholinesterase | T682A | 72 | A:0.917 | G:0.083 | Zhao et al. 2014 |
| supercont3.196 | 232328 | CPIJ008566 | CYP6Z15: cytochrome P450 6Z15 | E243A | 70 | A:0.857 | C:0.143 | Kothera et al. 2019 |
| supercont3.228 | 585169 | CPIJ009085 | CYP6AG13: cytochrome P450 6AG13 | N211D | 52 | A:0.827 | G:0.173 | Kothera et al. 2019 |
| supercont3.510 | 164957 | CPIJ014218 | CYP9M10: cytochrome P450 9M10 | F245I | 66 | A:0.864 | T:0.136 | Kothera et al. 2019 |

## 4 DISCUSSION

Here we have shown that these baits are a relatively cost-effective way to survey a sufficient number of segregating genetic sites taken from across the genomes of a large number of *Culex pipiens* complex samples. Specifically, we used our developed gene-based capture assay to successfully examine taxonomic relationships, population structure, and patterns of admixture in these mosquitoes. We also showed these baits can be used to survey the presence and frequency of known insecticide resistance alleles.

Perhaps not surprisingly, the genetic reads derived from *Cx. quinquefasciatus* samples mapped the best to the genome, while the outgroup sample, *Cx. torrentium*, mapped the most poorly (Table S3). When we just use the *Cx. quinquefasciatus* samples to look at the relationship between number of raw reads generated and the number of successfully mapped reads, we observed a small but significant, positive trend (Figure S3). This suggests that multiplexing fewer samples in a lane (e.g. MiSeq) may be preferable to increase the number of reads per sample, but there are likely other factors to consider. These may include the age of the sample (and corresponding DNA degradation), and the relative taxonomic distance from the reference. In the latter case, the number of variants which will be useful in downstream analyses may not be greatly improved by a greater depth of sequencing.

In our clustering analysis using principal components, we observed the greatest genetic distinction between the *Cx. quinquefasciatus* samples and those of the other taxa (Fig. 1). Interestingly, the samples of *Cx. quinquefasciatus* clustered more tightly than these other samples when considered collectively. This likely reflects the genetic divergence they also harbor. However, we also observed high levels of genetic diversity within these taxa, particularly *Cx. pipiens f. pipiens* and *Cx. pipiens pallens* (Table 1). It remains to be determined how much of this is true biological diversity, and how much could be an artifact of reference-based mapping biases. We also observed two primary genetic groups in our Admixture analysis (Fig. 2), with K=2 being the best supported (Figure S2). As with our PCA, this manifests as *Cx. quinquefasciatus* in one cluster and the other taxa in a second cluster. In both the PCA and Admixture analysis, the hybrid samples showed the expected mixture of lineages.

Interestingly, in our Admixture analysis at K=3, the *Cx. quinquefasciatus* samples became split between a Hawaiian and Samoan group and the rest of the samples. This is a surprising result given the patterns of clustering observed in the PCA, which differentiated *Cx. pipiens pallens* from the other taxa along the second axis. In the Admixture analysis, *Cx. pipiens pallens* only became distinct at K=5.

When we examined clustering in just the *Cx. quinquefasciatus* samples, we again observe the greatest differences between the Hawaiian and Samoan samples and everything else (Figures 1 & 3). However, for our Admixture analysis, K=1 was the best supported, which is not surprising given that these represent a single taxon with the potential for high rates of inter-population gene flow. Considering patterns of genetic diversity within *Cx. quinquefasciatus* populations, the east Asian samples harbored the highest mean number of heterozygous sites and a correspondingly high π value (Table 2). This recapitulates previous examinations of genetic diversity in this species (Fonseca et al. 2006). The lowest genetic diversity was present in the Brazilian samples, which may indicate a relatively recent colonization of South America.

In our quantitative examination of taxonomic differentiation, weighted F_st_ values were always higher than unweighted values (Table 3). Not surprisingly, the greatest F_st_ values observed were between the taxa in the species complex and the outgroup, *Cx. torrentium* (Table 3). Interestingly, among taxa in the species complex, the highest unweighted value was observed between *Cx. pipiens f. molestus* and *Cx. pipiens pallens*, whereas for weighted values it was between *Cx. pipiens f. molestus* and *Cx. quinquefasciatus*. The distinctiveness of the *Cx. Pipiens f. molestus* samples from these two taxa is also observed in our principal component analysis (Fig. 1). As expected, the lowest weighted and unweighted F_st_ values are both for the comparison of the two forms of *Cx. pipiens*.

Within *Cx. quinquefasciatus*, the greatest genetic differentiation was between the samples from Nigeria and those from Samoa (Table 4). This may reflect their relative geographic distance and the corresponding decrease in genetic exchange. However, other factors such as differential selection could also play a role in generating the genetic divergence observed between African and Samoan population of *Cx. quinquefasciatus*.

In our taxonomic analysis, the outgroup sample *Cx. torrentium* was distinct from the other samples, as expected (Fig. 4). Within the samples of the *Cx. pipiens* complex, there are two major clades comprised of the *Cx. quinquefasciatus* samples and everything else. The *Cx. quinquefasciatus* clade was well supported (99/100 bootstrap support), whereas the second clade was more poorly supported (52/100 bootstrap support), likely the effect of the hybrid samples which were all collected in North America. Interestingly, the samples of *Cx. pipiens* f. *pipiens* and *Cx. pipiens* f. *molestus* do not form monophyletic clades in this analysis. This may reflect the low level of genetic differentiation between the two taxa, combined with extensive genetic exchange between them.

Our assessment of insecticide resistance alleles revealed the presence of all identified variants in at least one of the sequenced samples. This points to the ubiquity and maintenance of these resistant alleles in the *Cx. pipiens* complex, and underscores the importance of careful insecticide resistance management (Dusfour et al. 2020). However, all but one of these variants were segregating at low frequencies (< 20% of the samples), which suggests that there are likely fitness costs to harboring these variants. There are known fitness costs associated with mutations in the acetylcholinesterase gene (Rivero et al 2011) in the absence of strong selection from insecticide exposure, and these may be generalizable to cytochrome P450 mutations, except possibly for those at 6BZ2.

In conclusion, the described bait-based assay is a powerful tool to address phylogenomic questions at multiple scales, including taxonomic differentiation and population structure across the *Cx. pipiens* complex. It can also be used to uncover the presence and extent of gene flow among populations and admixture. Furthermore, the utility of the data that can be generated using these baits is likely to expand. For example, it will be possible to investigate specific evolutionary drivers of taxonomic differentiation such as drift or selection. Of particular interest will be the identification of variation in specific genes contributing to the extensive ecological and behavioral differences observed among the *Cx. pipiens* complex taxa.

## ACKNOWLEDGEMENTS

We thank Alison Devault at Arbor Biosciences for assistance with bait design. This work was funded in part by NSF EAGER Award 1547168 and NSF Ecology and Evolution of Infectious Diseases Award 2001213 (PI DMF).

## Supplementary Tables

**Table S1**. Genes used in Bait Design

**Table S2**. Sample Information

**Table S3**. Sample sequencing and mapping stats

**Table S4**. Genetic diversity observations by all samples.

**Table S5**. Genetic diversity observations for just the *Cx. quinquefasciatus* samples.

**Table S6**. Insecticide resistance alleles by sample

## Supp. Figures

**Figure S1**. Variant Quality Distributions

**Figure S2**. Admixture CV results. a) all complex samples b) just *Cx. quinquefasciatus* samples

**Figure S3**. Correlation between number of raw reads sequenced and the percent of these reads that subsequently mapped to the reference genome for the *Cx. quinquefasciatus* samples.

