## Supplementary figures and images for "A gene-based capture assay for surveying patterns of genetic diversity and insecticide resistance in a worldwide group of invasive mosquitoes"

### Supplemental Figure 1

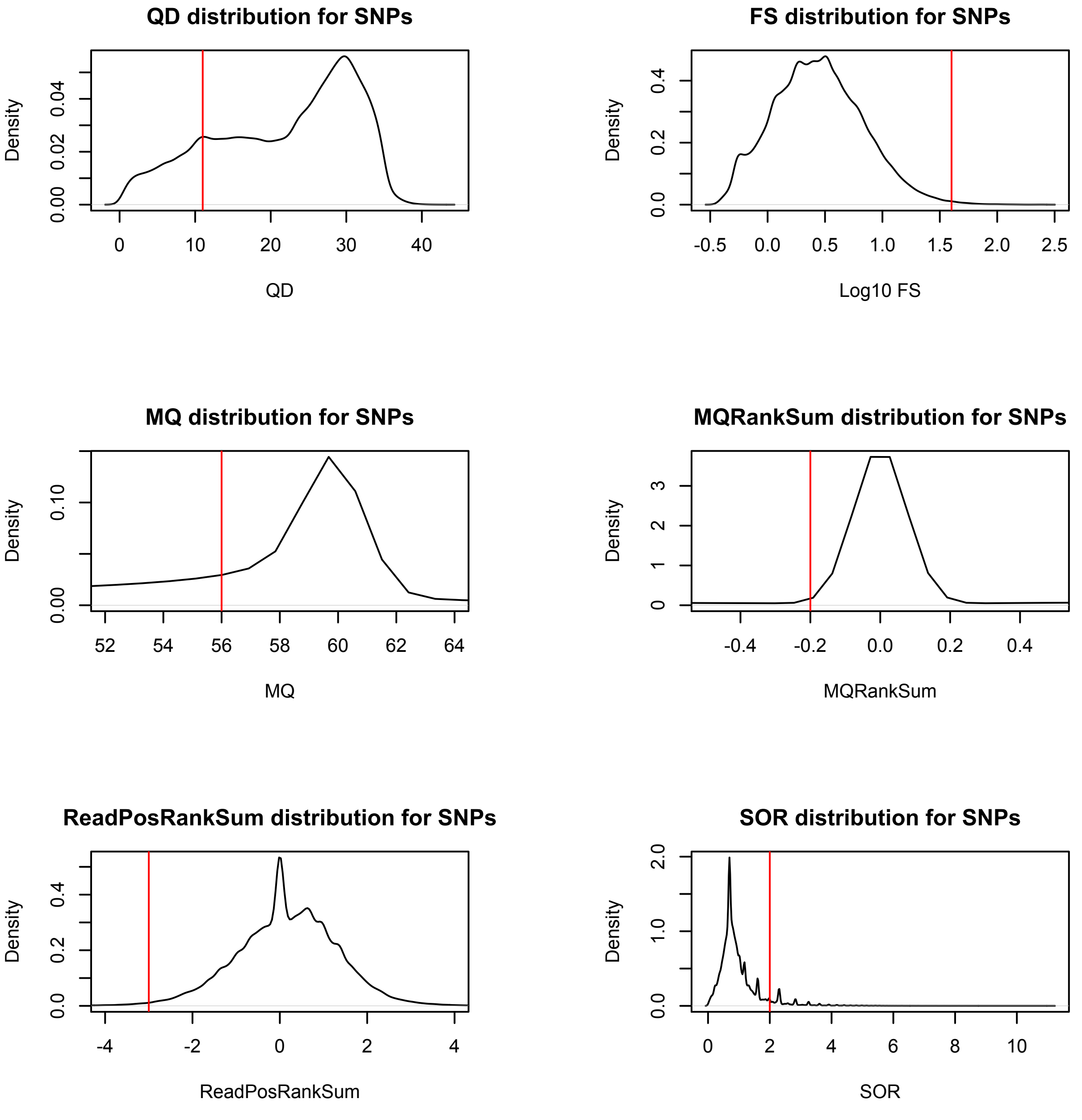

### Supplemental Figure 2

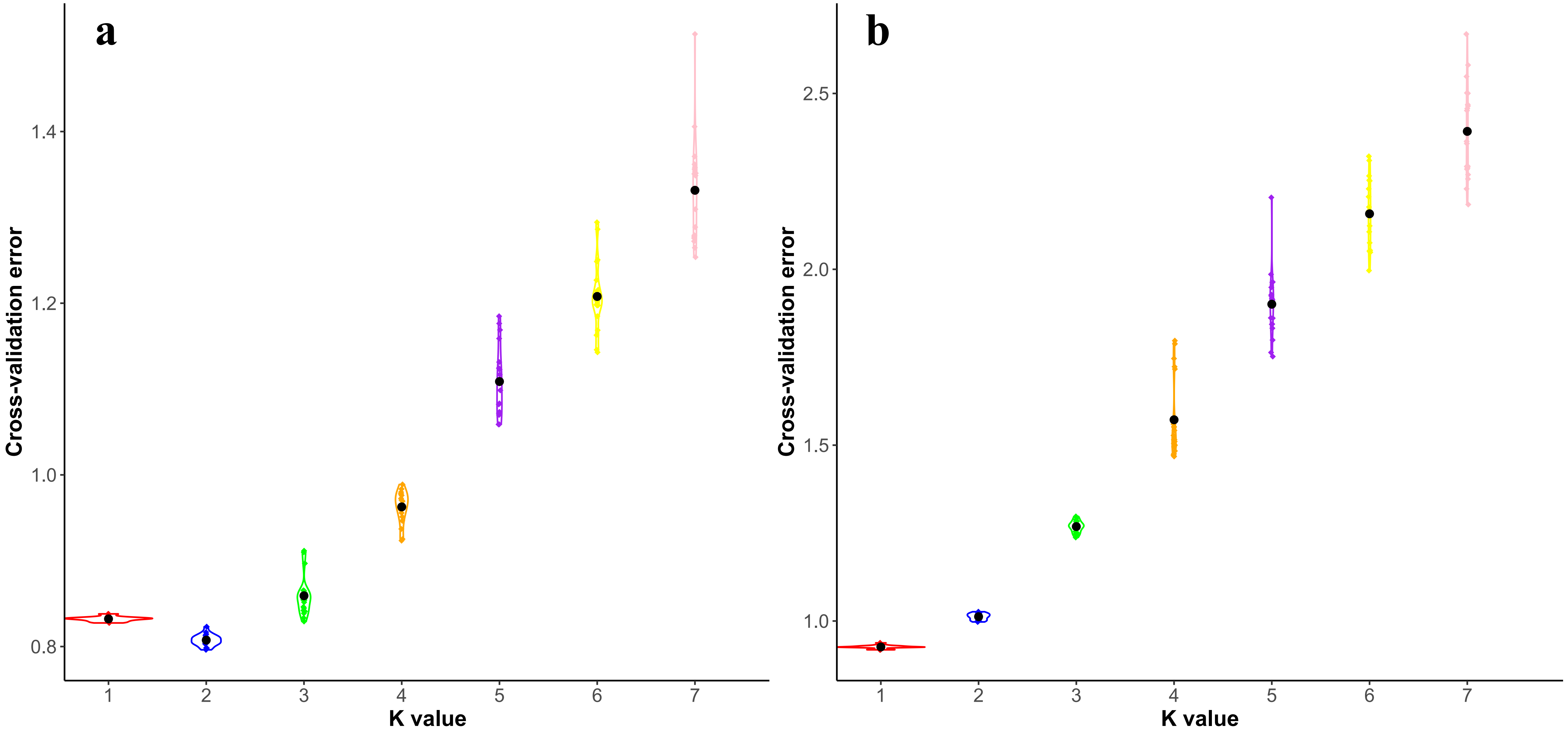


All Complex Samples

Only *Cx. quinquefasciatus* Samples

### Supplemental Figure 3

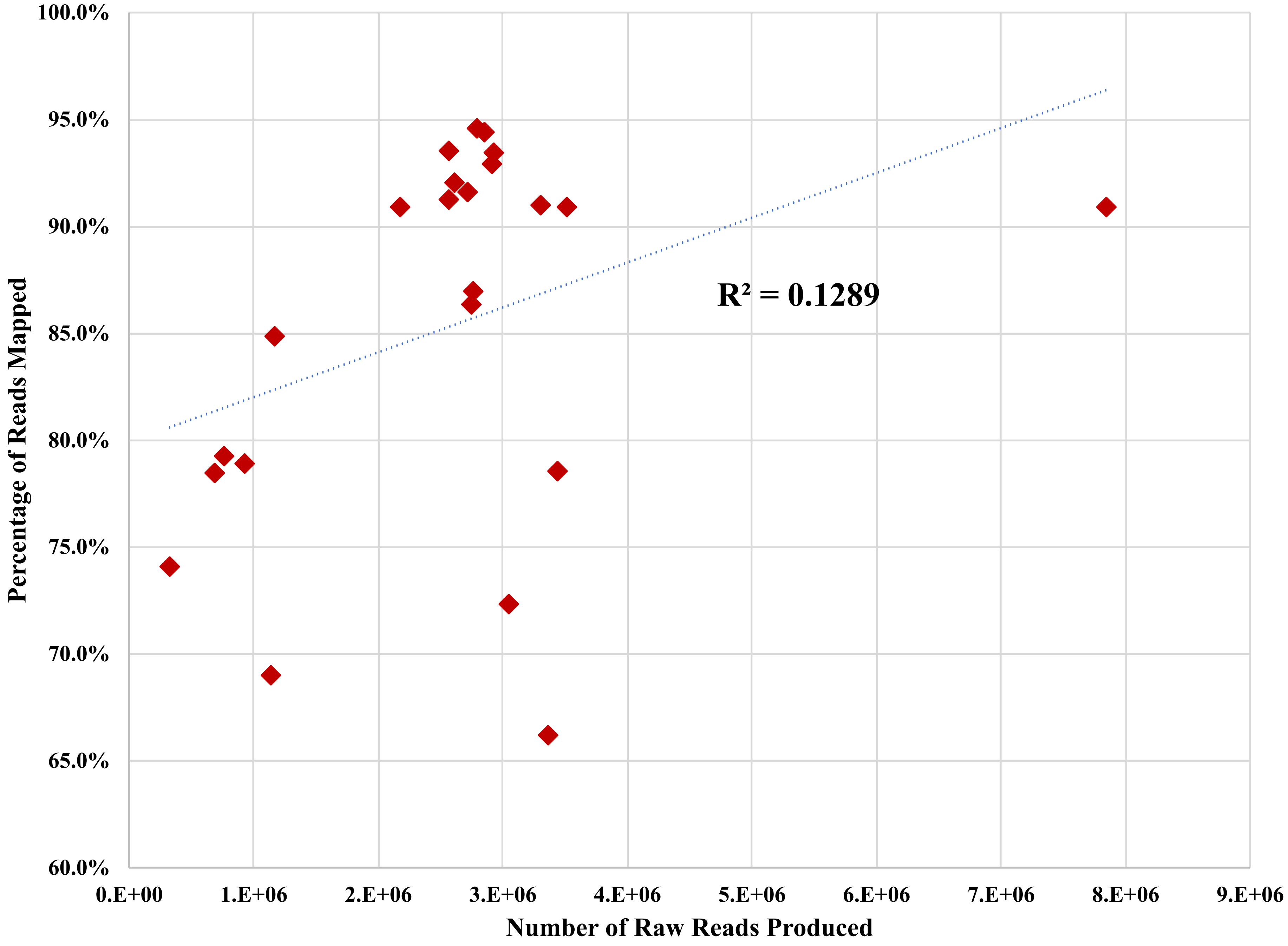
